# StomataCounter: a neural network for automatic stomata identification and counting

**DOI:** 10.1101/327494

**Authors:** Karl C. Fetter, Sven Eberhardt, Rich S. Barclay, Scott Wing, Stephen R. Keller

**Affiliations:** Department of Plant Biology, University of Vermont; Department of Paleobiology, Smithsonian Institution, National Museum of Natural History; Amazon.com, Inc

**Keywords:** Stomata, Computer Vision, Convolutional Neural Network, Deep Learning, Phenotyping

## Abstract

- Stomata regulate important physiological processes in plants and are often phenotyped by researchers in diverse fields of plant biology. Currently, there are no user friendly, fully-automated methods to perform the task of identifying and counting stomata, and stomata density is generally estimated by manually counting stomata.
- We introduce StomataCounter, an automated stomata counting system using a deep convolutional neural network to identify stomata in a variety of different microscopic images. We use a human-in-the-loop approach to train and refine a neural network on a taxonomically diverse collection of microscopic images.
- Our network achieves 98.1% identification accuracy on Ginkgo SEM micrographs, and 94.2% transfer accuracy when tested on untrained species.
- To facilitate adoption of the method, we provide the method in a publicly available website at http://www.stomata.science/.

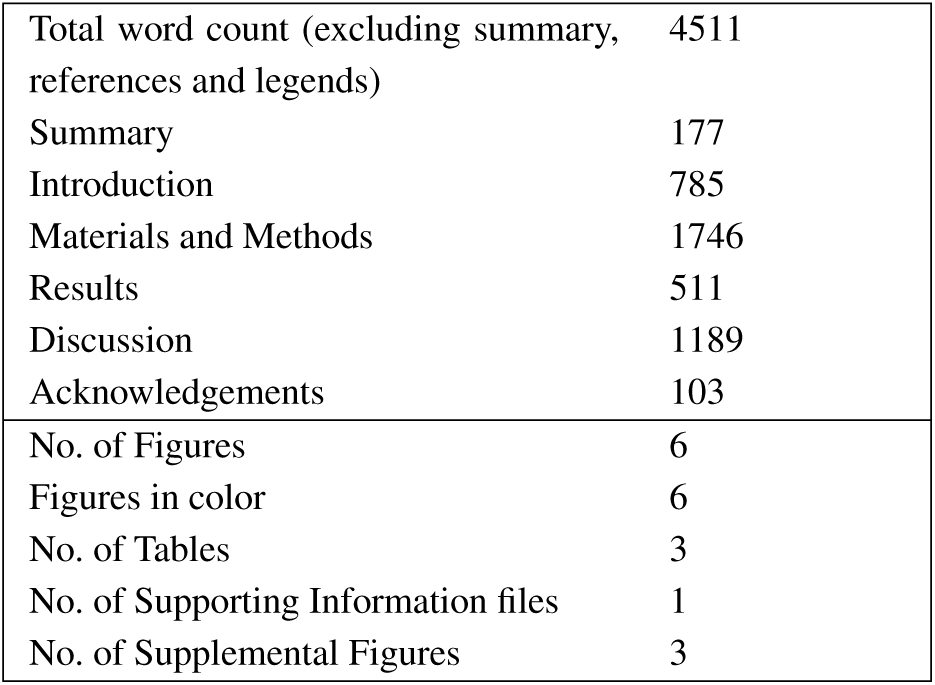

## I. INTRODUCTION

Stomata are important microscopic organs which mediate important biological and ecological processes and function to exchange gas with the environment. A stoma is composed of a pair of guard cells forming an aperture that, in some species, are flanked by a pair of subsidiary cells (Bergmann and Sack, 2007). Regulation of the aperture pore size, and hence, the opening and closing of the pore, is achieved through changing turgor pressure in the guard cells (Shimazaki *et al.*, 2007). Stomata evolved to permit exchange between internal and external sources of gases, most notably CO_2_ and water vapor, through the impervious layer of the cuticle (Kim *et al.*, 2010).

Because of their importance in regulating plant productivity and response to the environment, stomata have been one of the key functional traits of interest to researchers working across scales in plant biology. At the molecular level, regulation of stomata has been the subject of numerous genetic studies (see Shimazaki *et al.*, 2007 and (Kim *et al.*, 2010) for detailed reviews) and crop improvement programs have modified stomata phenotypes to increases yield (Fischer *et al.*, 1998). Stomata also mediate trade-offs between carbon gain and pathogen exposure that are of interest to plant ecophysiologists and pathologists. For example, foliar pathogens frequently exploit the aperture pore as a site of entry. In *Populus*, some species, and even populations within species, evolve growth strategies that maximize carbon fixation through increased stomatal density and aperture pore size on the adaxial leaf surface. This adaptation results in a cost of increased infection by fungal pathogens which have more sites of entry to the leaf (McKown *et al.*, 2014). Stomata, as sites of water vapor exchange, are also implicated in driving environmental change across biomes (Hetherington and Woodward, 2003) and variation of stomatal density and aperture pore length are linked to changes of ecosystem productivity (Wang *et al.*, 2015). Stomatal traits are of particular interest to paleoecologists and paleoclimatologists due to the relationship between stomatal traits and gas exchange. Measurements of stomatal density from fossil plants have been proposed as an indicator of paleoatmospheric CO_2_ concentration (Royer, 2001), and measuring stomatal traits to predict paleoclimates has become widely adopted (JC McElwain and Steinthorsdottir, 2017).

It is clear from their physiological importance that researchers across a wide variety of disciplines in plant biology will phenotype stomatal traits for decades to come. A typical stomatal phenotyping pipeline consists of collecting plant tissue, creating a mounted tissue for imaging, imaging of the specimen, and manual phenotyping of a trait of interest. This last step can be the most laborious, costly, and time-consuming task, reducing the efficiency of the data acquisition and analysis pipeline. This is especially important in large-scale plant breeding and genome-wide association studies, where phenotyping has been recognized as the new data collection bottleneck, in comparison to the relative ease of generating large genome sequence datasets (Hudson, 2008).

Here, we seek to minimize the burden of high-throughput phenotyping of stomatal traits by introducing an automated method to identify and count stomata from micrographs. Although automated phenotyping methods using computer vision have been developed (Higaki *et al.*, 2014; Laga *et al.*, 2014; Duarte *et al.*, 2017; Jayakody *et al.*, 2017), these highly specialized approaches require feature engineering specific to a collection of images. These methods do not transfer well to images created with novel imaging and processing protocols. Additionally, tuning hand-crafted methods to work on a general set of conditions is cumbersome and often impossible to achieve. Recent applications of deep learning techniques to stomata counting have demonstrated success (Bhugra *et al.*, 2018; Aono *et al.*, 2019), but are not easily accessible to the public and use training data sets sampled from few taxa.

Deep convolutional neural networks (DCNN) circumvent specialized approaches by training the feature detector along with the classifier (LeCun *et al.*, 2015). The method has been widely successful on a range of computer vision problems in biology such as medical imaging (see Shen *et al.* (2017) for a review) or macroscopic plant phenotyping (Ubbens and Stavness, 2017). The main caveat of deep learning methods is that a large number of parameters have to be trained in the feature detector. Improvements in network structure (He et al., 2016) and training procedure (Simonyan and Zisserman, 2014) have helped training of large networks which incorporate gradient descent learning methods and prove to be surprisingly resilient against overfitting (Poggio *et al.*, 2017). Nevertheless, a large number of correctly annotated training images is still required to allow the optimizer to converge to a correct feature representation. Large labeled training sets like the ImageNet database exist (Jia Deng *et al.*, 2009), but for a highly specialized problems, such as stomata identification, publicly available datasets are not available at the scale required to train a typical DCNN.

We solve these problems by creating a large and taxonomically diverse training dataset of plant cuticle micrographs and by creating a network with a human-in-the-loop approach. Our development of this method is provided to the public as a web-based tool called StomataCounter which allows plant biologists to rapidly upload plant epidermal image datasets to pre-trained networks and then annotate stomata on cuticle images when desired. We apply this tool to a the training dataset and achieve robust identification and counts of stomata on a variety of angiosperm and pteridosperm taxa.

## II. MATERIALS AND METHODS

### A. BIOLOGICAL MATERIAL

Micrographs of plant cuticles were collected from four sources: the cuticle database (https://cuticledb.eesi.psu.edu/, Barclay, J McElwain, *et al.*, 2007); a Ginkgo common garden experiment (Barclay and Wing, 2016); a new intraspecific collection of balsam poplar (*Populus balsamifera*); and a new collection from living material at the Smithsonian, National Museum of Natural History (USNM) and the United States Botanic Garden (USBG) (Table 1). Specimens in the cuticle database collection were previously prepared by clearing and staining leaf tissue and then imaged. The entire collection of the cuticle database was downloaded on November 16, 2017. Downloaded images contained both the abaxial and adaxial cuticles in a single image, and were automatically separated with a custom bash script. Abaxial cuticle micrographs were discarded if no stomata were visible or if the image quality was so poor that no stomata could be visually identified by a human. Gingko micrographs were prepared by chemically separating the upper and lower cuticles and imaging with an environmental SEM Barclay and Wing, 2016. To create the poplar dataset, dormant cuttings of *P. balsamifera* genotypes were collected across the eastern and central portions of its range in the United States and Canada by S.R. Keller *et al.* and planted in common gardens in Vermont and Saskatchewan in 2014. Fresh leaves were sampled from June to July 2015 and immediately placed in plastic bags and then a cooler for temporary storage, up to three hours. In a laboratory, nail polish (Sally Hansen, big kwik dry top coat #42494) was applied to a 1 cm^2^ region of the adaxial and abaxial leaf surfaces, away from the mid-vein, and allowed to dry for approximately twenty minutes. The dried cast of the epidermal surface was lifted with clear tape and mounted onto a glass slide. The collection at the USNM and USBG was made similarly from opportunistically sampled tissues of the gardens around the National Museum of Natural History building. USBG collections were focused on the Hawaiian, southern exposure, and medicinal plant, and tropical collections. The taxonomic identity of each specimen was recorded according to the existing label next to each plant. The USNM/USBG collection was imaged with an Olympus BX-63 microscope using differential interference contrast (DIC). Some specimens had substantial relief and z-stacked images were created using cellSens image stacking software (Olympus Corporation). Slides were imaged in three non-adjacent areas per slide. The poplar collection was imaged with an Olympus BX-60 using DIC and each slide was imaged in two areas. Mounted material for the USNM/USBG and Ginkgo collections are deposited in the Smithsonian Institution, National Museum of Natural History in the Department of Paleobiology. Material for the poplar collection is deposited in the Keller laboratory in the Department of Plant Biology at the University of Vermont, USA. The training dataset totaled 4618 images (Table 1).

**Table 1.** Description of the datasets used for training and testing the network.

| Dataset | Training |  | Test |  | Preparation method | Imaging method | Magnification | Z-stacks applied | Citation |
| --- | --- | --- | --- | --- | --- | --- | --- | --- | --- |
| | $N_{\text{images}}$ | $N_{\text{species}}$ | $N_{\text{images}}$ | $N_{\text{species}}$ | | | | | |
| Poplar | 3123 | 1 | 175 | 1 | Nail polish | DIC | 400 | no | This paper |
| Ginkgo | 408 | 1 | 200 | 1 | Lamina peel | SEM | 200 | no | Barclay and Wing, 2016 |
| Cuticle DB | 678 | 613 | 696 | 599 | Clear & stain | Brightfield | 400 | no | Barclay, J McElwain, *et al.*, 2007 |
| USNM/USBG | 409 | 124 | 696 | 132 | Nail polish | DIC, SEM<sup>1</sup> | 100<sup>2</sup>, 200, 400 | yes | This paper |
| Aorta | - | - | 116 | 1 | EVG + trichrome | Brightfield | 40 | no | Johnson and Cipolla, 2017 |
| Breast Cancer | - | - | 58 | 1 | Hematoxylin & eosin stain | Brightfield |  | no | Gelasca *et al.*, 2008 |
| Total | 4618 | 739 | 1941 | 735 |  |  |  |  |  |

Note: 1) Training SEM $N = 15$ ; 2) training set, $N = 4$ .

### B. DEEP CONVOLUTIONAL NEURAL NETWORK

We used a deep convolutional neural network (DCNN) to generate a stomata likelihood map for each input image, followed by thresholding and peak detection to localize and count stomata (Fig. 1). Because dense per-pixel annotations of stoma versus non-stoma are difficult to acquire in large quantity, we trained a simple image classification DCNN based on the AlexNet structure instead (Krizhevsky *et al.*, 2012), and copied the weights into a fully convolutional network to allow per-location assessment of stomata likelihood. Although this method does not provide dense per-pixel annotations, the resulting resolution proved to be high enough to differentiate and count individual stomata.

**Figure 1.**
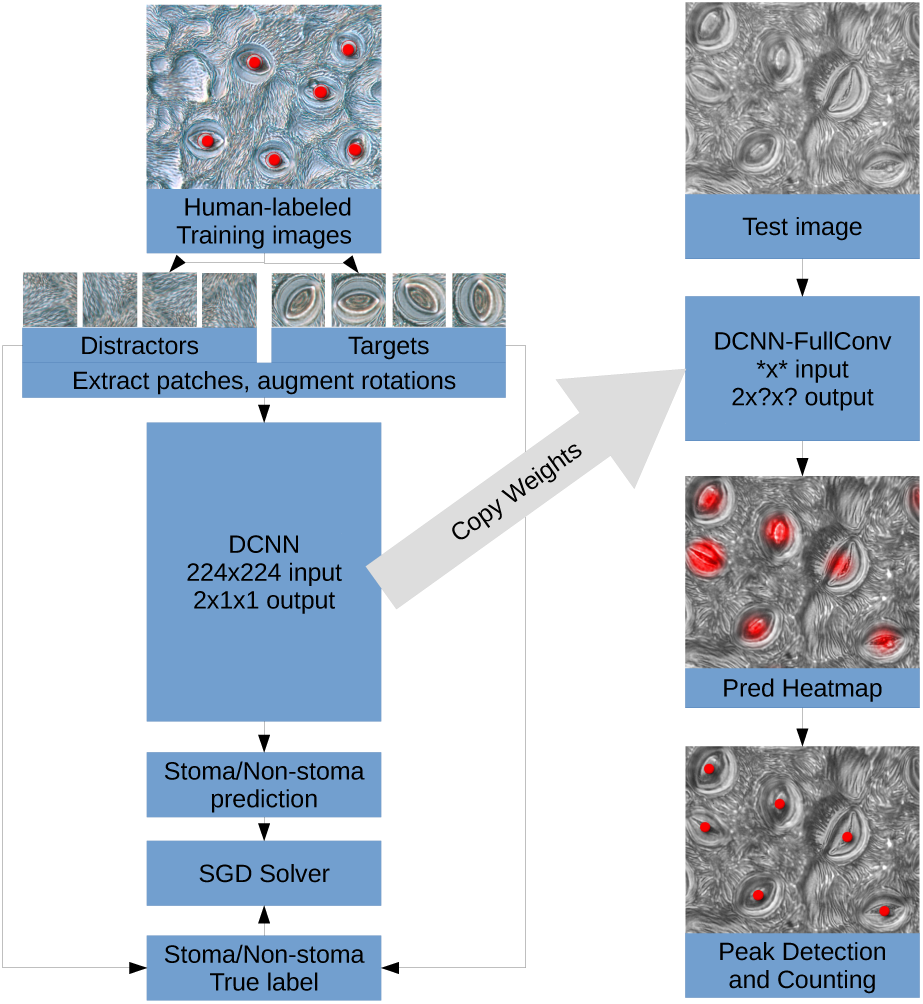
Left: Training and testing procedure. First column: Target patches were extracted and centered around human-labeled stomata center positions; distractor patches were extracted in all other regions. A binary image classification network was trained. Second column: The image classification network was applied fully convolutional to the test image to produce a prediction heatmap. On the thresholded heatmap, peaks were detected and counted.

We used pre-trained weights for the lowest five convolutional layers from conv1 to conv5. The weights were taken from the ILSVRC image classification tasks (Russakovsky *et al.*, 2015) made available in the caffenet distribution (Jia *et al.*, 2014). All other layers were initialized using Gaussian initialization with scale 0.01.

Training was performed using a standard stochastic gradient descent solver as in Krizhevsky *et al.* (2012), with learning rate 0.001 for pre-trained layers and 0.01 for randomly initialized layers with a momentum of 0.9. Because the orientation of any individual stomata does not hold information for identification, we augmented data by rotating all training images into eight different orientations, applied random flipping and randomly positioned crop regions of the input size within the extracted 256×256 image patch. For distractors, we sampled patches from random image regions on human-annotated images that were at least 256 pixels distant from any labeled stoma. The trained network weights were transferred into a fully convolutional network (Long *et al.*, 2015), which replaces the final fully connected layers by convolutions. To increase the resolution of the detector slightly, we reduced the stride of layer *pool*5 from two to one, and added a dilation (Yu and Koltun, 2015) of two to layer *fc*6_*conv*_ to compensate. Due to margins (96 pixels) and stride (32 over all layers), application of the fully convolutional network to an image of size s yields an output probability map *p* of size *s -*(96 *\** 2)*/*32 along each dimension.

To avoid detecting low-probability stomata within the noise, the probability map *p* was thresholded and all values below the threshold *p*-thresh=0.98 were set to zero. Local peak detection was run on a 3×3 pixel neighborhood on the thresholded map, and each peak, excluding those located on a 96-pixel width border, was labeled as a stoma center. This intentionally excludes stomata for which the detection peak is found near the border within the model margin to match the instructions given to human annotators. Resulting stomata positions were projected back onto the original image (see figure 2).

**Figure 2.**
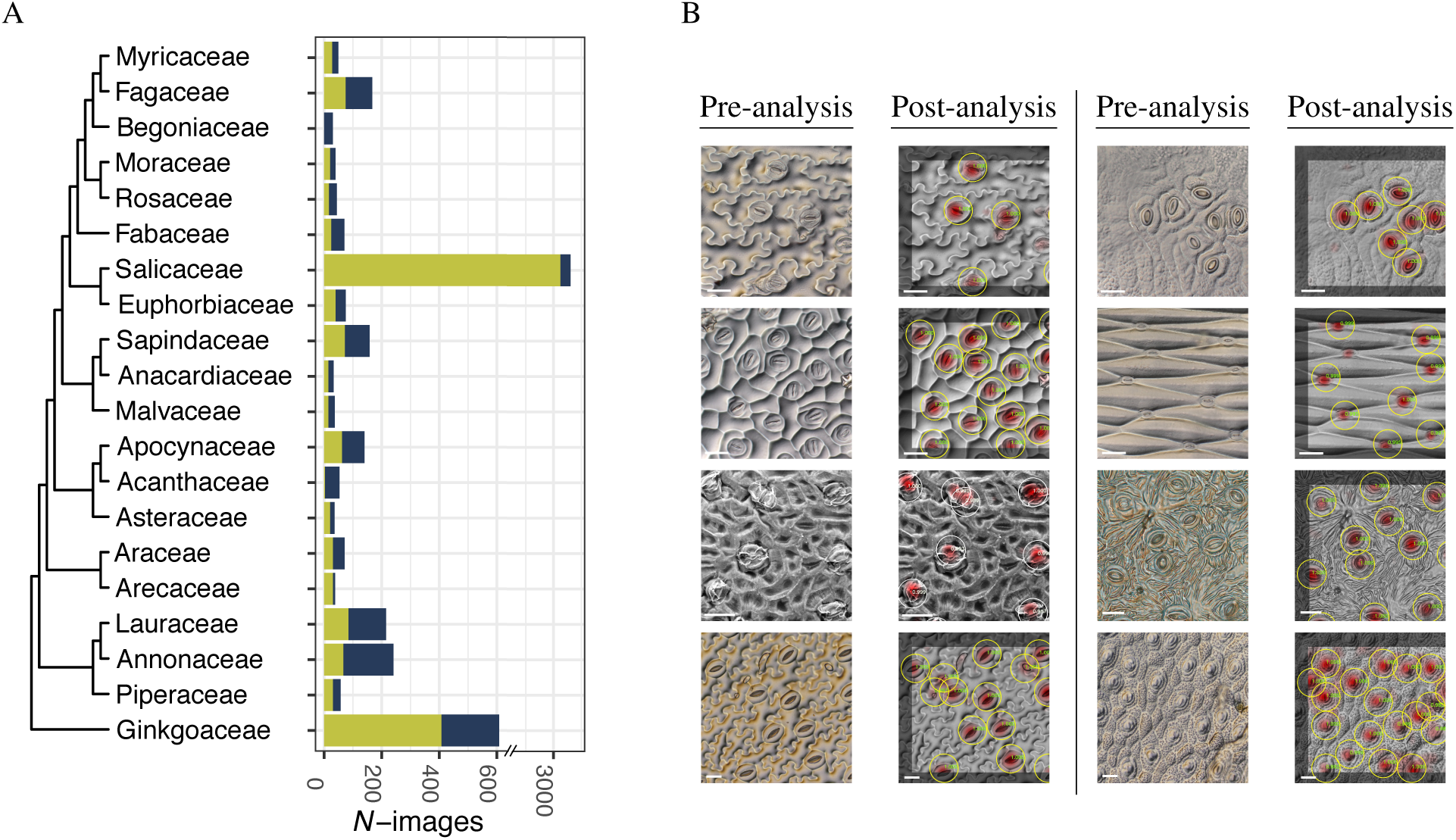
Sample sizes of images for the top 20 families represented in the training and test dataset (A). Examples of pre- and post-analysis images. A probability heatmap map is overlain onto the input image in the red channel. Detected stomata marked with circles with peak values given in green (B). Species in image pairs: *Adiantum peruvianum, Begonia loranthoides* (top row); *Chaemaerauthum gadacardii, Echeandia texensis* (second row); *Ginkgo biloba, Populus balsamifera* (third row); *Trillium luteum, Pilea libanensis* (bottom row). Scale bars in row one and two equal 20 *µ*m, scale bars in row three equal 25 *µ*m, and scale bars in row four equal 50 *µ*m.

We built a user-friendly web service, StomataCounter, freely available at http://stomata.science/, to allow the scientific community easy access to the method. We are using a flask/jquery/bootstrap stack. Source codes for network training, as well as the webserver are available at http://stomata.science/source. To use StomataCounter, users upload single .jpg images or a zip files of of their .jpg images containing leaf cuticles prepared. Z-stacked image sets should be combined into single image before uploading. A new dataset is then created where the output of the automatic counts, image quality scores and image metadata are recorded and can easily be exported for further analysis.

In addition to automatic processing, the user can manually annotate stomata and determine the empirical error rate of the automatic counts through a straightforward and intuitive web interface. The annotations can then be reincorporated into the training dataset to improve future performance of StomataCounter by contacting the authors and requesting retraining of the DCNN.

### C. STATISTICAL ANALYSES

We tested the performance of the DCNN with a partitioned set of images from each dataset source. Whenever possible, images from a given species were used in either the training or test set, but not both, and only 7 out of 1467 species are included in both. In total, 1941 images were used to test the performance of the network (Table 1). After running the test set through the network, stomata were manually annotated. If the center of a stoma intersected the bounding box around the perimeter of the image, it was not counted.

To evaluate the DCNN, we first determined if it could identify stomata when they are known to be present and fail to identify them when they are absent. To execute this test, a set of 25 randomly selected abaxial plant cuticle micrographs containing stomata were chosen from each of the four datasets for a total of 100 images. To create a set of test images known to lack stomata, 100 adaxial cuticle micrographs were randomly sampled from the cuticle database. Visual inspection confirmed that none of the adaxial images contained stomata. Micrographs of thoracic aorta from an experimental rat model of preeclampsia (Johnson and Cipolla, 2017), and breast cancer tissue micrographs (Gelasca *et al.*, 2008) were used as negative controls from non-plant material.

Identification accuracy is tested by applying the DCNN to a small image patch either centered on a stoma (target), or taken from at least 256 pixels distance to any labeled stomata (distractor). This yields true positive (*N*_*TP*_), true negative (*N*_*TN*_), false positive (*N*_*FP*_) and false negative (*N*_*FN*_) samples. We define the classification accuracy *A* as:

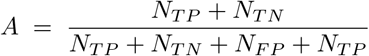

We define classification precision *P* as:

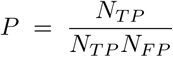

We assume the human counts contain only true positives and the automatic count contain true positives and false positives. As such, we define precision as:

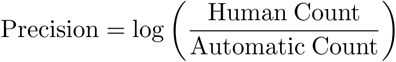

This definition of precision identifies over-counting errors as negative values and under-counting as positive values. This measure of precision is undefined if either manual or automatic count is zero, and 30 of the 1772 observations were discarded. These samples were either out of focus, lacked stomata entirely, or too grainy for human detection of stomata. Classification recall *R* was defined as:

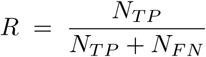

Classification accuracy, precision, and recall, were calculated from groups of images constructed to span the diversity of imaging capture methods (i.e. brightfield, DIC, and SEM) and magnification (i.e. 200x and 400x) to determine how well the training set from one group transfers to identifying stomata in another group. Groups used for transfer assessment were the cuticle database, the Ginkgo collection, micrographs imaged at 200x, at 400x, and the combined set of images.

We use linear regression to understand the relationship between human and automatic stomata counts. For the purpose of calculating error statistics, we consider deviation from the human count attributable to error in the method. Images were partitioned for linear models by collection source, higher taxonomic group (i.e. Eudicot, Monocot, Magnoliid, Gymnosperm, Fern, or Lycophyte), and magnification. To understand how different sources of variance contribute to precision variance, we collected data on the taxonomic family, magnification, imaging technique, and three measures of image quality. The taxonomic family of each image was determined using the open tree of life taxonomy accessed with the rotl package (Michonneau *et al.*, 2016). Two image quality measures used were based on the properties of an image’s power-frequency distribution tail and described the standard deviation (fSTD) and mean (fMean) of the tail. Low values of fSTD and fMean indicate a blurry image, while high values indicate non-blurry images. The third image quality measure, entropy, is a measure an image’s information content. High entropy values indicate high contrast/noisy images while low values indicate lower contrast. These image quality measures were created with PyImq (Koho *et al.*, 2016) and standardized between zero and one. Random effects linear models were created with the R package lme4 (Bates *et al.*, 2014) by fitting log precision to taxonomic family, magnification, and imaging technique as factors. Linear models were fit for the scaled error and image quality scores. We used the root mean square error (RMSE) of the model residuals to understand how the factors and quality scores described the variance of log precision. Higher values of RMSE indicate larger residuals. Statistical analyses were conducted in python and R (R Team, 2016). Count results and metadata for the test and training set available in supplementary table 1 and 2, respectively. Images used for the training and test sets are available as zip files (place holder for dryad upload).

## III. RESULTS

### A. STOMATA DETECTION

StomataCounter was able to accurately identify and count stomata when they were present in an image; stomata were detected in all of the abaxial cuticle images. False positives were detected in the adaxial cuticle, aorta, and breast cancer cell image sets at low frequency (Fig. S1). The mean number of stomata detected in the adaxial, aorta, and breast cancer image sets was 1.5, 1.4, and 2.4, respectively, while the mean value of the abaxial set containing stomta was 24.1.

Correspondence between automated and human stomata counting varied among the respective sample sets. There was close agreement among all datasets to the human count, with the exception of some of the samples imaged at 200x magnification (Fig. 3A-C). In these samples, the network tended to under count relative to human observers. Despite the variation among datasets and the large error present in the 200x dataset, the slopes of all models were close to 1 (Table 2). The 400x, Ginkgo, and cuticle database sets all performed well at lower stomata counts, as indicated by their proximity to the expected one-to-one line and decreased in precision as counts increased.

**Table 2.** Summary of linear model fit parameters in Figure 3 for different test datasets. Dataset definitions given in text. Abbreviations: y-intercept, *a*; standard error, *SE*. †200x images removed from this USNM/USBG set. Significance indicated by: *: *P* < 0:05; **: *P* < 0:01; ***: *P* < 0:001.

| (A) Dataset |  |  |  |  |  |
| --- | --- | --- | --- | --- | --- |
|  | Cuticle DB | Poplar | Ginkgo | USNM/USBG | USNM/USBG† |
| $a$ | 3.315*** | 0.823 | -0.364 | 10.068*** | -0.695 |
| $SE_a$ | (0.635) | (0.456) | (0.677) | (2.041) | (0.5) |
| $s$ | 0.853*** | 0.978*** | 0.832*** | 0.997*** | 1.167*** |
| $SE_s$ | (0.029) | (0.014) | (0.028) | (0.089) | (0.026) |
| $r^2$ | 0.578 | 0.964 | 0.812 | 0.157 | 0.835 |

| (B) Taxonomic group |  |  |  |  |  |  |
| --- | --- | --- | --- | --- | --- | --- |
|  | Eudicot | Monocot | Magnoliid | Gymnosperm | Fern | Lycophyte |
| $a$ | 10.405*** | 2.360*** | 4.360* | 0.004 | 0.624 | 0.771 |
| $SE_a$ | (1.643) | (0.598) | (1.844) | (0.735) | (0.474) | (1.414) |
| $s$ | 0.808*** | 0.808*** | 0.912*** | 0.830*** | 0.799*** | 0.751** |
| $SE_s$ | (0.063) | (0.056) | (0.086) | (0.031) | (0.039) | (0.169) |
| $r^2$ | 0.141 | 0.617 | 0.272 | 0.778 | 0.938 | 0.739 |

| (C) Magnification |  |  |
| --- | --- | --- |
|  | 200x | 400x |
| $a$ | 18.346*** | 1.446*** |
| $SE_a$ | (3.192) | (0.381) |
| $s$ | 0.654*** | 0.970*** |
| $SE_s$ | (0.123) | (0.017) |
| $r^2$ | 0.055 | 0.740 |

**Figure 3.**
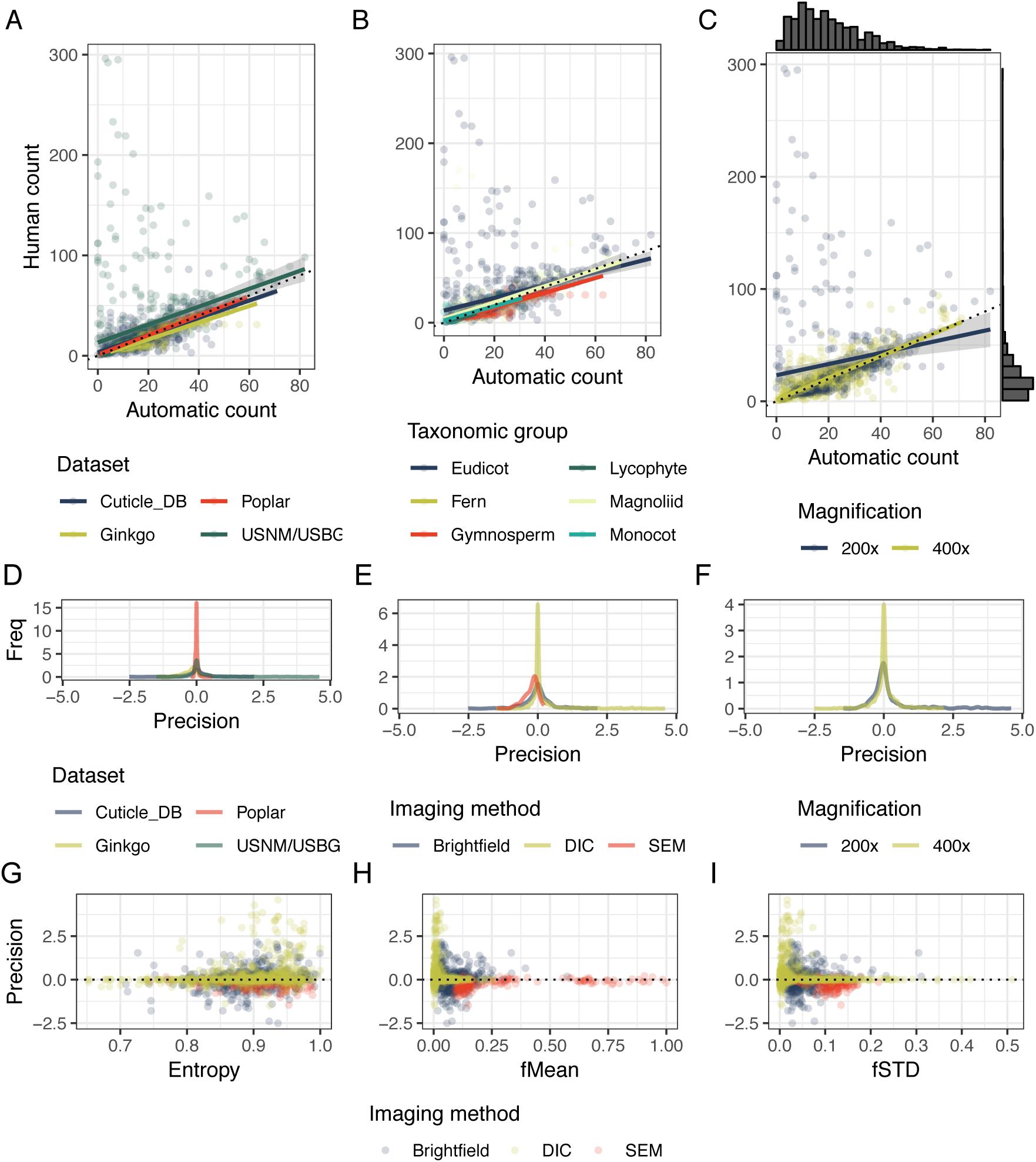
Results of human and automatic counts and precision for each dataset organized by collection source (A,D), taxonomic group (B), and magnification (C), and imaging method (E). Precision and image quality scores have non-linear relationships, and changes in variance correlate to image quality (G-I). Positive values of precision indicate undercounting of stomata relative to human counts, while negative values indicate overcounting. The dotted black line is the 1:1 line, indicating where perfect automatic counts occur in a plot. See Table 2 for model summaries.

Precision has a non-linear relationship with image quality, and changes in variance of precision are correlated with variation of image quality. With a ratio of 17:1 images in the training vs. test sets, the Poplar dataset had the best precision, followed by the Ginkgo, Cuticle database, and USNM/USBG datasets (Fig. 3D). Among the different imaging methods, SEM had the best precision, followed by DIC, and finally and brightfield microscopy (Fig. 3E). The variance of precision is higher for images with 200x magnification (Fig. 3F). RMSE values were lowest for taxonomic family and the family:magnification interaction, suggesting these factors contributed less to deviations between human and automated stomata counts than image quality or imaging method (Table 3).

**Table 3.** Root mean square error (RMSE) of the model residuals. Lower RMSE values suggest a better fit of the model. The response in each model was precision.

| Model | RMSE |
| --- | --- |
| ~ family | 0.463 |
| ~ magnification | 0.523 |
| ~ family:magnification | 0.399 |
| ~ image_technique | 0.519 |
| ~ fMean | 0.523 |
| ~ fSTD | 0.523 |
| ~ entropy | 0.521 |
| ~ entropy:imaging_method | 0.515 |

### B. CLASSIFICATION ACCURACY

The peak accuracy (94.2%) on the combined test sets is achieved when all training sets are combined 5. The combined dataset performs best on all test subsets of the data; i. e. adding additional training data - even from different sets - is always beneficial for the generalization of the network. Accuracy from train to test within a single species is higher (e. g. Ginkgo training for Ginkgo test at 97.4%) than transfer within datasets with a large number species across families (400x training to 400x test: 85.5%).

The network does not generalize well between vastly different scales, i.e. the 200x-dataset, which contains images downscaled to half the image width and height. In this case, only training within the same scale achieves high accuracy (97.3%), while adding additional samples from the larger scale reduces the performance (to 90.7%).

Precision values are generally higher than recall (0.99 precision on the combined training and test sets; 0.93 recall, see supplement 2), which shows that we mostly miss stomata rather than misidentifying non-existing stomata.

Increased training size is correlated with increased accuracy 6, and providing a large number of annotated images is beneficial, as it lifts training accuracy from 72.8% with a training set of ten images to 94.2% with the complete set of training images.

## IV. DISCUSSION

Stomata are an important functional trait to many fields within plant biology, yet manual phenotyping of stomata counts is a laborious method that has few controls on human error and reproducibility. We created a fully automatic method for counting stomata that is both highly sensitive and reproducible, allows the user to quantity error in their counts, but is also entirely free of parameter optimization from the user. Furthermore, the DCNN can be iteratively retrained with new images to improve performance and adjust to the needs of the community. This is a particular advantage of this method for adjusting to new taxonomic sample sets. However, new users are not required to upload new image types or images from new species for the method to work on their material. The pre-loaded neural network was specifically trained on diverse set of image types and from many species (N = 739).

As the complexity of processing pipelines in biological studies increases, repeatability of studies increasingly becomes a concern (Bruna, 2014, Open Science Foundation, 2015). Apart from the reduced workload, automated image processing provides better reproducibility than manual stomata annotations. For instance, if multiple experts count stomata, they may not agree, causing artificial differences between compared populations. This includes how stomata at the edge of an image are counted, and what to do with difficult to identify edge cases. Automatic counters will have an objective measure, and introduce no systematic bias between compared sets as long as the same model is used. Additionally, our human counter missed stomata that the machine detected (e.g. Fig. 4).

**Figure 4.**
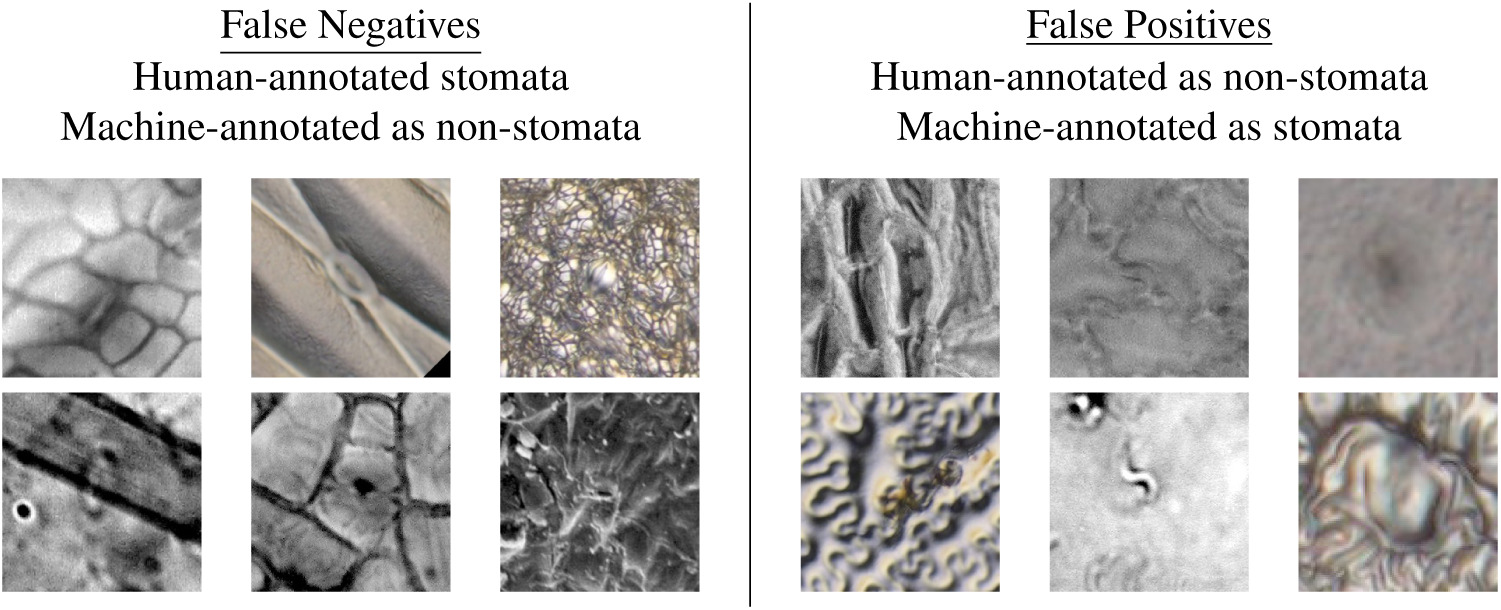
Samples that were mislabeled with high confidence by either the machine or human. False negatives (left panel) from stomata that were detected by the human, but not by the machine. Image features typically generating false negatives are blurry images, artifacts obstruction the stomata, low contrast between epidermal and guard cells, and very small scale stomata. False positives (right panel) are image features that are labeled by the machine as stomata, but not by the human counter, or are missed by the human counter (e.g. top and bottom right images in the panel). Errors typically generating false positives are image artifacts that superficially resemble stomata, particularly shapes mimicking interior guard cell structure.

Our method is not the first to identify and count stomata. However, previous methods have not been widely adopted by the community and a survey of recent literature indicates manual counting is the predominant method (Liu *et al.*, 2018; Morales-Navarro *et al.*, 2018; Sumathi *et al.*, 2018; Peel *et al.*, 2017; Takahashi *et al.*, 2015). Previous methods relied on substantial image pre-processing to generate images for thresholding to isolate stomata for counting (Oliviera *et al.*, 2014; Duarte *et al.*, 2017). Thresholding can perform well in a homogenous collection of images, but quickly fails when images collected by different preparation and micrscopy methods are provided to the thresholding method (K. Fetter, pers. obs.). Some methods also require the user to manually segment stomata and subsequently process those images to generate sample views to supply to template matching methods (Laga *et al.*, 2014). Object-oriented methods (Jian *et al.*, 2011) also require input from the user to define model parameters. These methods invariably requires the user to participate in the counting process to tune parameters and monitor the image processing, and are not fully autonomous. More recently, cascade classifier methods have been developed which perform well on small collections of test sets (Vialet-Chabrand and Brendel, 2014; Jayakody *et al.*, 2017). Additionally, most methods rely on a very small set of images (50 to 500) typically sampled from just a few species or cultivars to create the training and/or test set (e.g., Bhugra *et al.*, 2018).

Apart from generalization concerns, several published methods require the user to have some experience coding in python or C++, a requirement likely to reduce the potential pool of end users. Our method resolves these issues by being publicly available, fully autonomous of the user, who is only required to upload jpeg formatted images, is free of any requirement for the user to code, and is trained on a relatively large and taxonomically diverse set of cuticle images.

We have demonstrated that this method is capable of accurately identifying stomata when they are present, but false positives may still be generated by shapes in images that approximate the size and shape of stomata guard cells. Conversely, false negatives are generated when a stomata is hidden by a feature of the cuticle or if poor sample preparation/imaging introduces blur. This issue is likely avoidable through increased sample size of the training image set and good sample preparation and microscopy techniques by the end user.

The importance of having a well-matched training and testing image set was apparent at 200x, where there was a subset of observations where transfer accuracy was low (Fig. 5), and StomataCounter consistently under counted relative to human observation (Fig. 3). We argue that transferring architecture between scales is not advisable and images should be created by the user to match the predominant size (2048×2048) and magnification (400x) of images in the training network. Our training set of images spanned 82 different families and was over-represented by angiosperms. Stomata in gymnosperms are typically sunken into pores that make it difficult to obtain good nail polish casts. Models tested to explain variation in scaled error revealed that taxonomic family and its interaction with magnification were the factors that had the best explanatory power for scaled error. Future collections of gymnosperm cuticles could be uploaded to the DCNN to retrain it in order to improve the performance of the method for gymnosperms.

**Figure 5.**
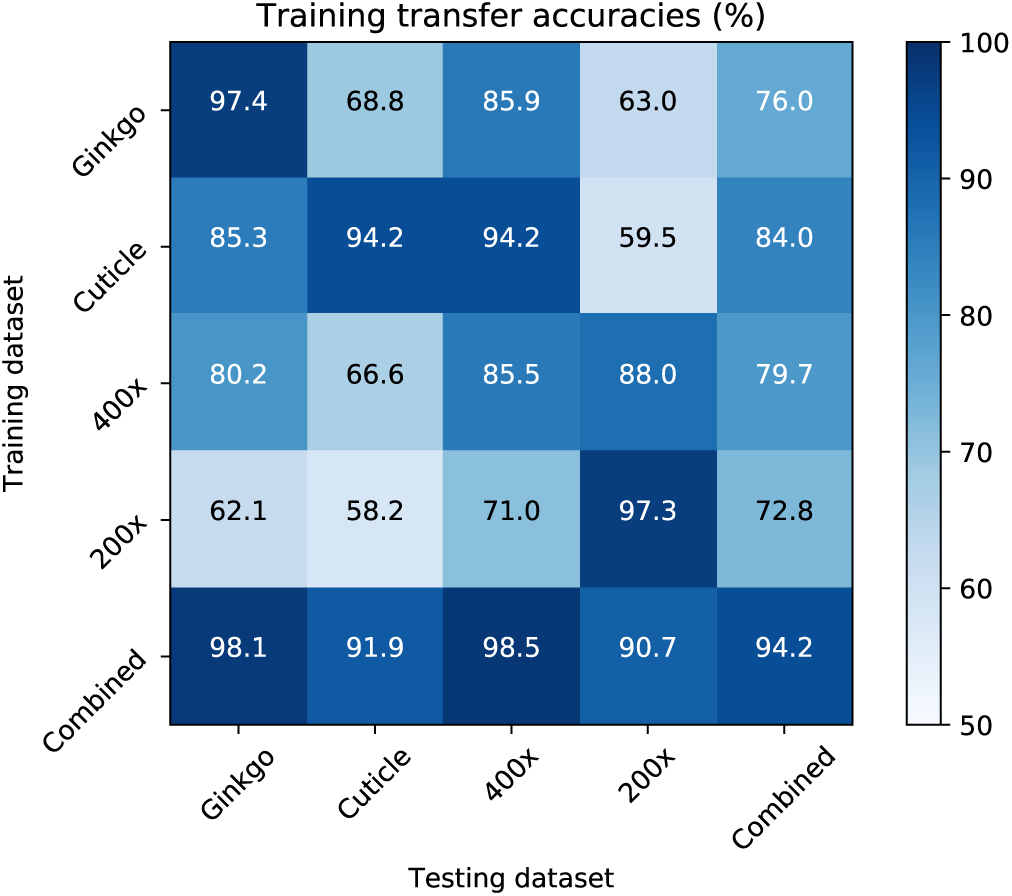
Accuracies for models trained on different training datasets (vertical), tested on different test datasets. Combined is a union of all training and test sets. For precision and recall values, see supplement 2.

**Figure 6.**
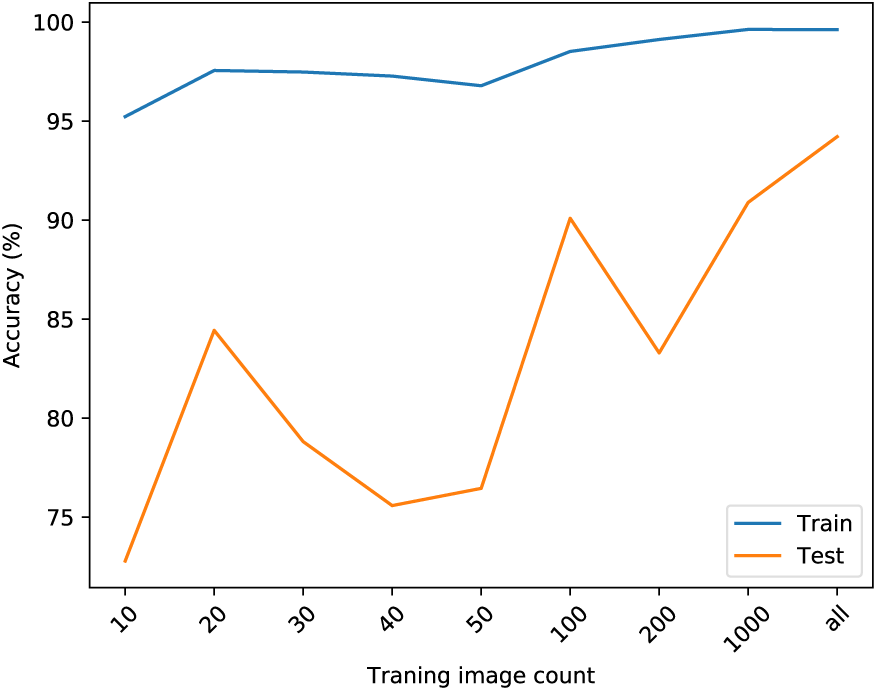
Accuracy by training image count and the effect of increasing the training set size on classification accuracies. Since classification is a binary task, chance level is at 50%. Training image count “*all*” includes all 4,618 annotated training images.

More generally however, this highlights how users will need to be thoughtful about matching of training and test samples for taxa that may deviate in stomata morphology from the existing reference database. We therefore recommend that users working with new or morphologically divergent taxa first run several pilot tests with different magnification and sample preparation techniques to find optimal choices that minimize error for their particular study system. SEM micrographs had the least amount of error, followed by DIC, and finally brightfield (Fig. 3G-I). Lastly, image quality was strongly related to log precision; predictably, images that are too noisy (i.e. high entropy) and out-of-focus (low fMean or fSTD) will generate higher error. Obtaining high quality, in-focus images should be a priority during data acquisition. We provide these guidelines for using the method, and recommend that users read these guidelines before collecting a large quantity of images:

- Collect sample images using different microscopy methods from the same tissues. We recommend an initial collection of 25 to 100 images prior to initiating a new large-scale study.
- Run images through StomataCounter.
- Establish a ‘true’ stomata count using the annotation feature.
- Regress image quality scores (automatically provided in output csv file) against log precision.
- Regress human versus automatic counts and assess error.
- Choose the microscopy method that minimizes error and image the remaining samples.

Different microscopy methods can include using DIC or phase contrast filters, adjusting the aperture to increase contrast, or staining tissue and imaging with under fluorescent light (see Eisele *et al.*, 2016 for more suggestions). If a large collection of images is already available and re-imaging is not feasible, we recommend the users take the following actions:

- Randomly sample 100 images.
- Upload the images to StomataCounter.
- Annotate images to establish the ‘true’ count.
- Explore image quality scores with against the log(precision) to determine a justifiable cut-off value for filtering images.
- Discard images below the image quality cut-off value.

New users can also contact the authors through the StomataCounter web interface and we may retrain the model to include the 100 annotated images.

Fast and accurate counting of stomata increases productivity of workers and decreases the time from collecting a tissue to analyzing the data. Until now, assessing measurement error required phenotyping a reduced set of images multiple times by, potentially, multiple counters. With StomataCounter, users can instantly phenotype their images and annotate them to create empirical error rates. The open source code and flexibility of using new and customized training sets will make StomataCounter and important resource for the plant biology community.

## V. ACKNOWLEDGEMENTS

We thank Kyle Wallick at the United States Botanic Garden for facilitating access to the living collections of the Garden. Scott Whitaker aided in imaging cuticle specimens with SEM and DIC microscopy in the Laboratories of Analytical Biology at the Smithsonian Institution, National Museum of Natural History. Rat aorta images were provided by Dr. A. Chapman and Dr. M. Cipolla at the University of Vermont. Dr. Terry Delaney at the University of Vermont kindly allowed KCF to use his microscope for imaging. Funding to create Stomata Counter was provided by an NSF grant to SRK (IOS-1461868) and a Smithsonian Institution Fellowship to KCF.

## VI. DATA AVAILABILITY

Zip files of the training and test images will be uploaded to dryad (This is a placeholder/reminder).

## VII. AUTHOR CONTRIBUTIONS

The research was conceived and performed by KCF and SE. The website and python scripts were written by SE. Data were collected and analyzed by KCF. Ginkgo Images were submitted by RSB and SW. All authors interpreted the results. The manuscript was written by KCF, SE, and SRK. All authors edited and approved the manuscript.

## SUPPLEMENTAL FIGURES

**Supplemental Figure 1.**
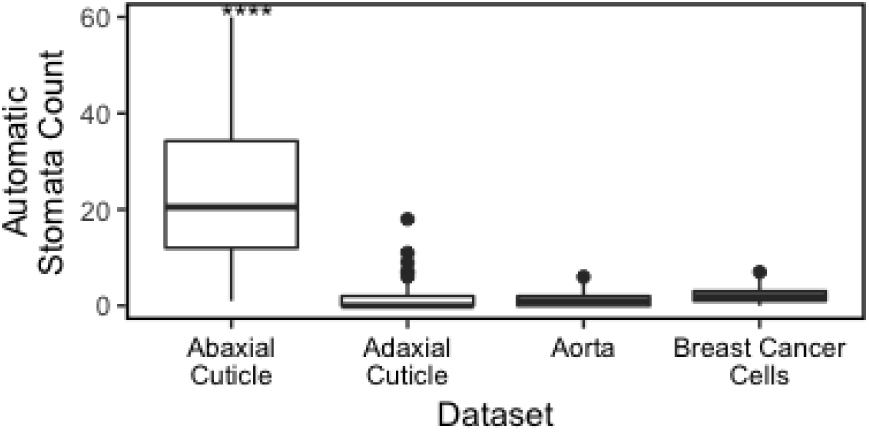
StomataCounter correctly identified stomata in all abaxial cuticle images containing stomata, while producing a low frequency of false positive counts in other image types. More false positives were identified in adaxial cuticle images than in images from human aorta or breast cancer cells. Resultx003Cs of post-hoc significance testing of pairwise means indicated by: *: *P* < 0.05; **: *P* < 0.01; ***: *P* < 0.001.

**Supplemental Figure 2.**
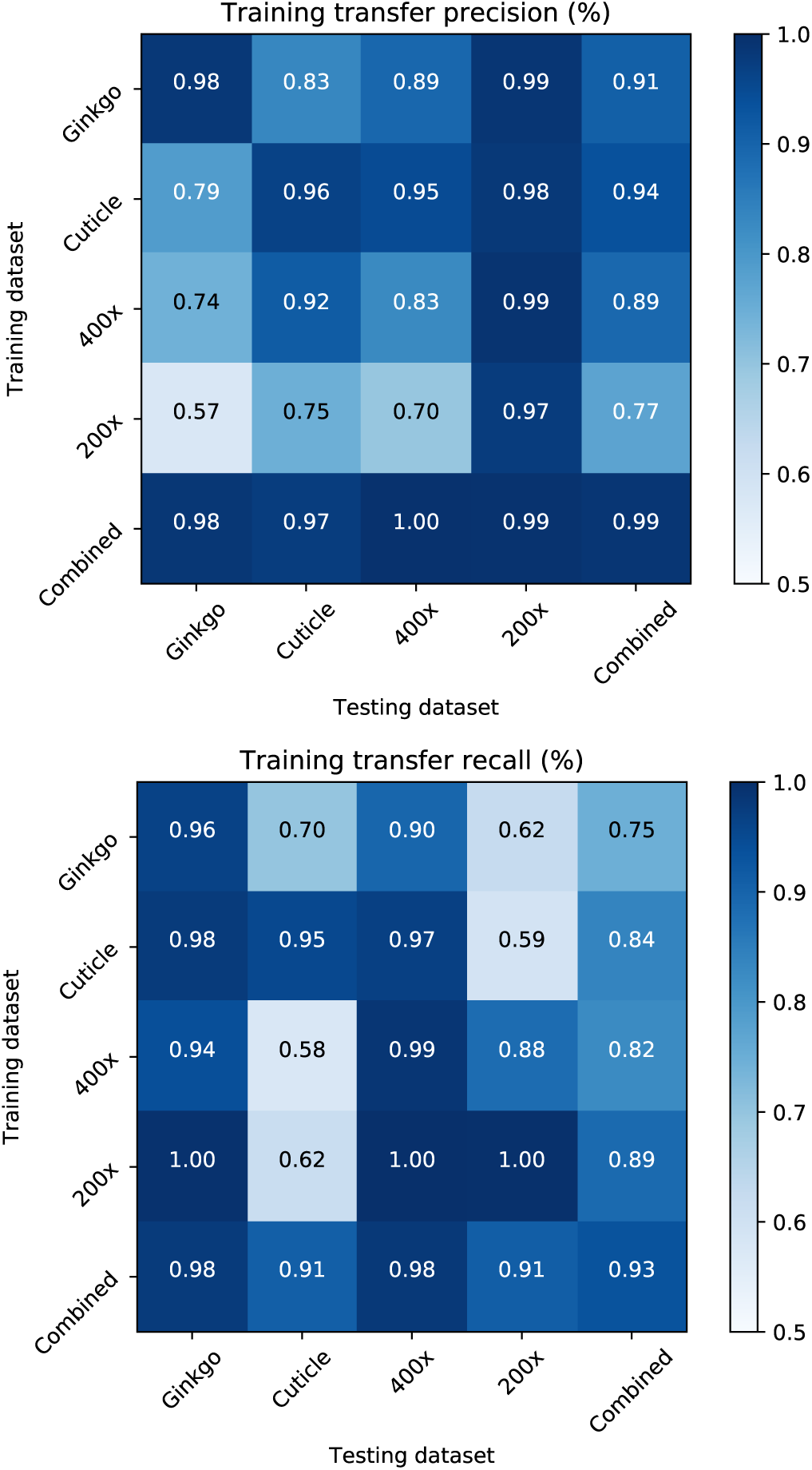
Precision and Recall for models trained on different training datasets (vertical), tested on different test datasets. *Combined* is a union of all training and test sets.

